# Self-Documenting Plasmids

**DOI:** 10.1101/2024.10.29.620927

**Authors:** Sarah I. Hernandez, Samuel J. Peccoud, Casey-Tyler Berezin, Jean Peccoud

## Abstract

Plasmids are the workhorse of biotechnology. These small DNA molecules are used to produce recombinant proteins and to engineer living organisms. They can be regarded as the blueprints of many biotechnology products. It is, therefore, critical to ensure that the sequences of these DNA molecules match their intended designs. Yet, plasmid verification remains challenging. To secure the exchange of plasmids in research and development workflows, we have developed self-documenting plasmids that encode information about themselves in their own DNA molecules. Users of self-documenting plasmids can retrieve critical information about the plasmid without prior knowledge of the plasmid identity. The insertion of documentation in the plasmid sequence does not adversely affect their propagation in bacteria and does not compromise protein expression in mammalian cells. This technology simplifies plasmid verification, hardens supply chains, and has the potential to transform the protection of intellectual property in the life sciences.

## Introduction

The role of plasmids in research and commercial production cannot be understated. Plasmids can be designed to express any sequence and engineered into living organisms to produce proteins, vaccines, viral vectors, gene circuits, and more [1-5]. The accuracy of plasmid sequences is critical to their reliable production and proper function. Sequence verification throughout a plasmid’s lifecycle (i.e., production, sharing, propagation, experimentation) is the best defense against inevitable random mutations, errors in production, and other undesired changes [6]. Yet, sequence verification is only possible when high-quality documentation about a plasmid is available.

Although plasmids can be made and sequenced with base-level precision, there is a history of poor validation and documentation practices surrounding DNA [7-9]. Approaches using restriction enzymes, PCR, or functional analysis to validate plasmids remain popular despite no guarantees that sequences have not changed. Furthermore, it is not uncommon to only have access to a plasmid map showing the plasmid structure without providing its sequence, stymieing sequence verification. When plasmid sequences are provided, the ambiguity of sequence annotations can make it challenging to understand the plasmid function [10]. Plasmid developers have little incentive to document the DNA sequences they engineer. They cannot claim authorship because DNA sequences are not copyrightable [11,12] and the scientific publishing enterprise puts little pressure on authors to release usable plasmid documentation [13,14]. Novel documentation practices are needed to give plasmid authors the means to get credit for their creative works and control their distribution.

The lack of a suitable framework to claim authorship of plasmids and their documentation is one of the factors contributing to a widespread disconnect between the parts and plasmids submitted to and retrieved from public and commercial repositories [15-17]. The other complicating factor is the dual nature of DNA sequences that can exist in two digital and physical forms. It is challenging to ensure that physical samples match the digital sequences that define them [9] because the dissemination of plasmids eventually leads to the corruption of the plasmid molecule, its documentation, or both (Figure 1). In addition, the community needs a system to uniquely identify physical plasmids associated with digital documentation. A similar problem was faced in the automotive industry: how can documentation related to a specific car’s manufacturer, title, insurance, etc., be identified amongst many instances of the same make and model? The solution was developing the vehicle identification number (VIN) system, which assigns unique digital certificates encoding information about a car’s identity and source to each car to associate it with its digital documentation. We propose a similar system, the GenoFAB Standard Identification Number (GSIN), that provides a framework to connect individual instances of a plasmid sequence with their unique documentation [18].

**Figure 1:**
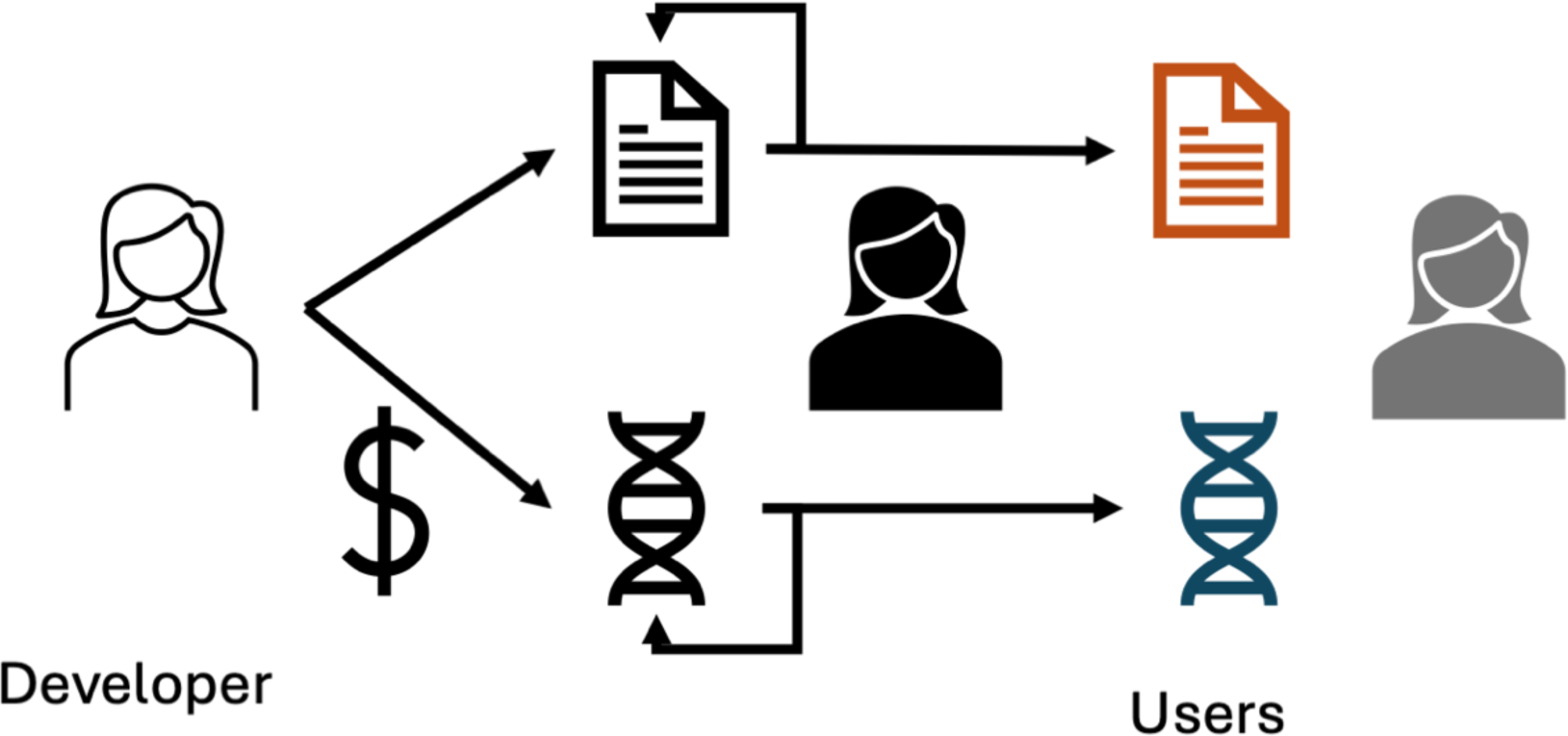
Plasmid sharing. Plasmids and their documentation (i.e., sequence, annotations, functional data, legal documents) are easy to make, copy, share, and change, accidentally or intentionally. Plasmid users may have computer files and DNA solutions that seem to be related because they share a common name. However, a similar name does not guarantee that the file and molecule match. Mutations accumulate when plasmids are used or shared. Plasmid may be edited without changing their name or documenting the changes. Similarly, many computer files used to document plasmids can be edited by the users without changing their names.

We leverage the data storage capabilities of DNA to achieve this goal. Cryptographic approaches like substitution ciphers (e.g., using “AGG” for “D”, “CTG” for “N”, and “GCA” for “A” to encode “DNA” in 9 nucleotides) enable DNA sequences to encode alphanumeric information like ID numbers, data files, and books [19-22]. This can be used to embed identifying information about a plasmid, or a reference to digital documentation [23], into its own sequence. These are often referred to as DNA watermarks or barcodes. Watermarked and barcoded DNA can be inserted into cells and is an easily stored and replicated data storage tool [24-27]. These approaches are not secure: the same watermark or barcode may be used on multiple documents, and it is possible to change, remove, or coopt watermarks and barcodes onto other documents [19]. In the physical world, security against identity or document counterfeits can be mitigated through a process like notarization. In the digital space, sophisticated cryptographic approaches known as digital signatures can provide similar assurances [19]. They mimic the security of an in-person notarization process by facilitating digital identity and documentation verification. Digital signatures use public key cryptography to allow the public verification of digital documents, their creators, and owners, as exemplified by digital services like DocuSign and recently popularized by non-fungible tokens (NFTs) [28].

We developed the first iteration of a digital certificate encoding minimal documentation directly in plasmid sequences [29-31]. This first-generation certificate relies on an RSA-based signature [32] to generate a digital signature encoded in 512 bp. The certificate, like a VIN, is composed of distinct data blocks encoding identifying information and security assurances, including a plasmid ID, author ID, and an error correction code to safeguard the signature and plasmid sequences. We validated this system using a handful of plasmids expressed in bacteria [31]. We also showed it was possible to use the Sakai-Kasahara scheme [33] to reduce the size of the signature to 164 bp. We also proposed to encode sequence annotations in the certificate. While we simulated the retrieval of a GenBank file using a FASTA file of the plasmid sequence, we had not experimentally validated these advancements.

Here, we experimentally validate the use of the 164 bp signature in combination with files embedded and stored within a plasmid to create self-documenting plasmids. We assess the feasibility of certifying a larger set of diverse plasmids and their use in both bacterial and mammalian cells. We show that plasmids can encode their own documentation files with little to no effect on the cells expressing them, and the files can be retrieved from *de novo* sequencing assemblies. We make the certification software available for plasmid developers and users to generate and verify certified plasmids. The certification of plasmids accelerates the verification process by eliminating the need for reference alignment, ensures plasmid users can assess the proper documentation, and provides security assurances in plasmid distribution.

## Results

### Generation of Self-Documenting Plasmids

We considered two different strategies to allow plasmid users to retrieve their plasmid documentation from sequencing data (Figure 2). First, we developed plasmids that provide their documentation as a link to a file residing on a server. These plasmids have a minimal certificate embedded and only provide a reference to their documentation (Figure 2a). We also developed plasmids that embed their documentation in their sequence without relying on computer files (Figure 2b,c).

**Figure 2:**
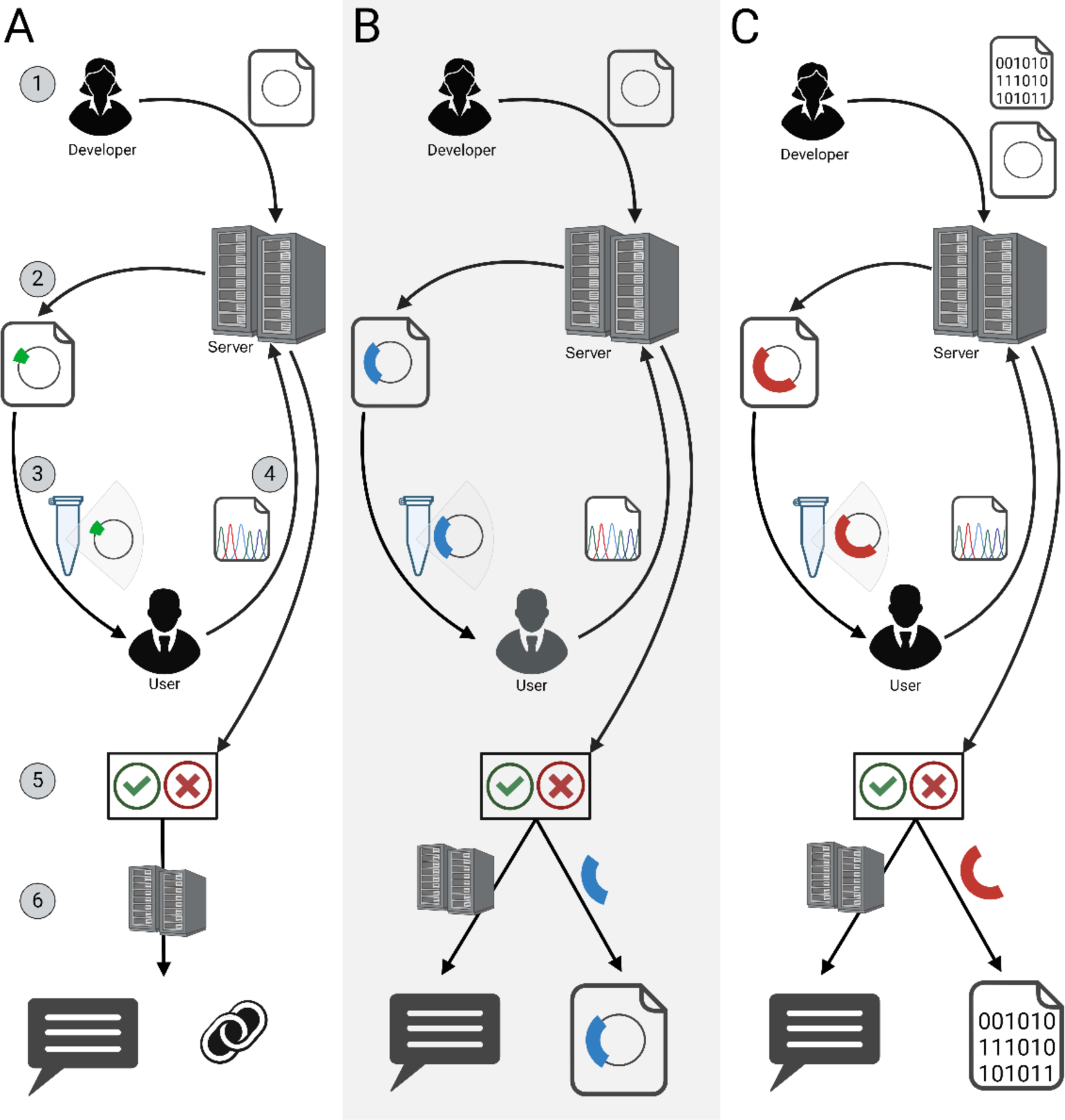
Certification workflow. Three types of certificates were generated and verified in similar workflows: (A) Certificate with 164 bp signature and reference to the documentation (green), (B) certificate with GenBank file embedded in sequence (blue), and (C) certificate with data file embedded in sequence (red). In each case, (1) the plasmid developer uploads a GenBank file containing an annotated plasmid sequence to the server. A backup of the GenBank file as text, plasmid ID, developer ID, and a URL to digital documentation are stored on the server. When applicable, data files to be embedded in the plasmid are also uploaded but not stored on the server (C). (2) Signature sequences and GenBank files for certified plasmids are generated. (3) Plasmids are synthesized and shared with plasmid users. (4) Users sequence the plasmids they received and upload the *de novo* assemblies to the server. (5) The signatures are compared against the information stored in the server and either pass or fail the verification process. (6) When successful, a report containing the plasmid ID and developer ID is retrieved from the server. The URL associated with the certified plasmid by the developer is also retrieved from the server (A). GenBank (B) and other files (C) embedded in the signature are retrieved directly from the sequence.

#### Documentation reference

We selected 24 plasmids of different origins to illustrate this self-documentation strategy (Table 1). Each of the plasmids has a descriptive name, *e.g.* CMV-mEGFP, and an alphanumerical identifier like DPLA3122. For each of these plasmids, we aimed to produce two functionally equivalent variants that could be distinguished by their serial number (Plasmid ID in Table 1). Starting from the 24 GenBank files describing the original plasmids, we generated 48 GenBank files corresponding to the 48 self-documenting plasmids (Figure S1). These self-documenting plasmids include a digital certificate in the form of a 260 bp long computer-generated DNA sequence that encodes the identity of the plasmid author, a unique plasmid serial number, and security features (Figure 3). In addition, each plasmid record is associated with a URL linking the plasmid to its documentation.

**Figure 3.**
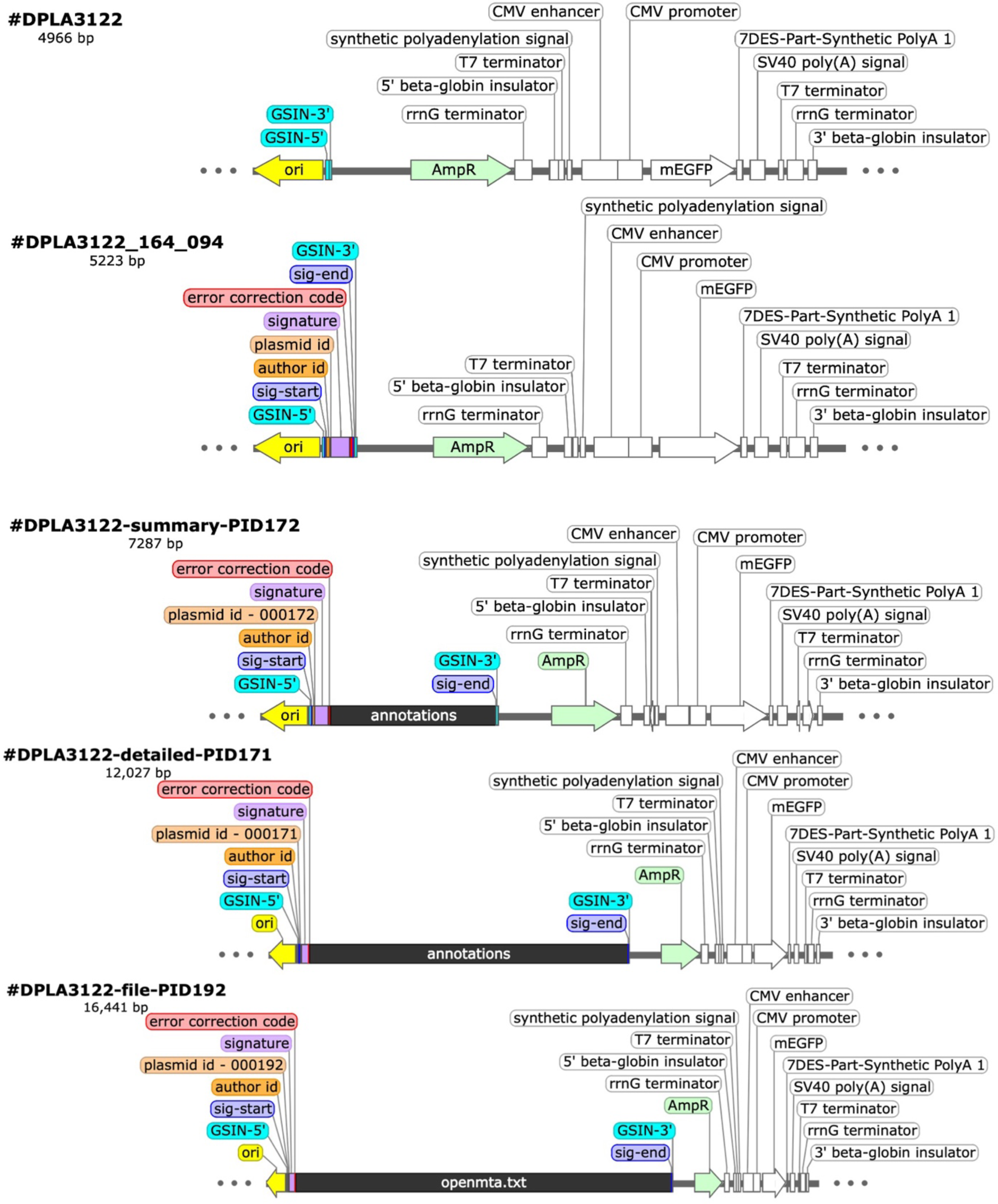
Structure of self-documenting plasmids. The same plasmid sequence (DPLA3122) was certified with all signature schemes (164 bp signature, summary GenBank, detailed GenBank, and openmta.txt file). GSIN 5’ and 3’ primers were used to linearize the plasmid and ligate the certificate into the plasmid backbone. The full certificate for the 164 bp signature is 260 bp and is comprised of distinct data blocks: the sig-start and sig-end delimiters used for certificate detection (10 bp each), author ID (32 bp), plasmid ID (12 bp), the signature containing the hashed (compressed) plasmid sequence for sequence verification (164 bp), and the error correction code (32 bp). GenBank files are encoded in an annotations feature, while text files are stored in a filename.txt feature, both of which scale with the size of the data file.

**Table 1:** Results of self-documenting plasmid assembly.

| Plasmid Name | Identifier | Plasmid ID | Verifies | Notes |
| --- | --- | --- | --- | --- |
| CMV-T7RNAP-Puro | DPLA3116 | 86 | Yes |  |
| CMV-T7RNAP-Puro | DPLA3116 | 87 | No, ECC fails | Large insertion, primer dimers |
| CMV-Int-mEGFP | DPLA3117 | 88 | Yes |  |
| CMV-Int-mEGFP | DPLA3117 | 89 | Yes |  |
| CMV-Int-mCherry2C | DPLA3118 | 90 | No, ECC fails | 2 bp insertion |
| CMV-Int-mCherry2C | DPLA3118 | 91 | Yes |  |
| CMV-Int-mEGFP-P2A-T7RNAP | DPLA3119 | 92 | Yes |  |
| CMV-Int-mEGFP-P2A-T7RNAP | DPLA3119 | 93 | No, ECC fails | Primer Dimers |
| CMV-mEGFP | DPLA3122 | 94 | No, ECC fails | Large insertion, primer dimers |
| CMV-mEGFP | DPLA3122 | 95 | Yes |  |
| pCMV-T7-EGFP | DPLA3125 | 96 | No, ECC fails | Large insertion, primer dimers |
| pCMV-T7-EGFP | DPLA3125 | 97 | Yes |  |
| pUC19-T7 pro-IRES-EGFP | DPLA3126 | 98 | No, ECC fails | 2 bp insertion |
| pUC19-T7 pro-IRES-EGFP | DPLA3126 | 99 | No, ECC fails | 2 bp insertion |
| pVSV-eGFP-dG | DPLA3135 | 100 | No, ECC fails | Signed, Missing VSV genes |
| pVSV-eGFP-dG | DPLA3135 | 101 | Yes, with ECC | 1 mismatch |
| CMV-mClover3 | DPLA3136 | 102 | Yes |  |
| CMV-mClover3 | DPLA3136 | 103 | Yes |  |
| CMV-DsRed2 | DPLA3137 | 104 | Yes, with ECC | 2 mismatches |
| CMV-DsRed2 | DPLA3137 | 105 | Yes |  |
| CMV-mCherry2C | DPLA3140 | 106 | No, ECC fails | 48 mismatch, 1 bp deletion |
| CMV-mCherry2C | DPLA3140 | 107 | Yes |  |
| CMV-moxEBFP | DPLA3142 | 108 | Yes |  |
| CMV-moxEBFP | DPLA3142 | 109 | No, ECC fails | 1 mismatch, 2 bp deletion |
| CMV-mScarlet-I | DPLA3146 | 112 | Yes |  |
| CMV-mScarlet-I | DPLA3146 | 113 | Yes |  |
| CMV-mTagBFP2 | DPLA3148 | 114 | No, ECC fails | Large insertion, primer dimers |
| CMV-mTagBFP2 | DPLA3148 | 115 | Yes |  |
| pVSVdG-4BFP2 | DPLA3152 | 116 | Yes |  |
| pVSVdG-4BFP2 | DPLA3152 | 117 | No, ECC fails | 5 mismatches, 16 indels |
| CMV-T7Pro-Int-mEGFP-P2A-T7RNAP | DPLA3181 | 118 | No, ECC fails | Large Deletion |
| CMV-T7Pro-Int-mEGFP-P2A-T7RNAP | DPLA3181 | 119 | No, ECC fails | Large Deletion |
| mCherry-mEGFP | DPLA3182 | 120 | No, ECC fails | 1 mismatch, 1 bp deletion |
| mCherry-mEGFP | DPLA3182 | 121 | Yes |  |
| pCMV-T7RNAP | DPLA3183 | 122 | No, ECC fails | Large insertion, primer dimers |
| pCMV-T7RNAP | DPLA3183 | 123 | Yes |  |
| pVSV-Venus-VSV-G | DPLA3187 | 124 | Yes |  |
| pVSV-Venus-VSV-G | DPLA3187 | 125 | No, ECC fails | Large insertion, primer dimers |
| CMV-VSV-P | DPLA3188 | 126 | Yes |  |
| CMV-VSV-P | DPLA3188 | 127 | Yes |  |
| CMV-VSV-N | DPLA3189 | 128 | No, ECC fails | Large insertion, primer dimers |
| CMV-VSV-N | DPLA3189 | 129 | No, ECC fails | Large insertion, primer dimers |
| CMV-VSV-M | DPLA3190 | 130 | Yes |  |
| CMV-VSV-M | DPLA3190 | 131 | Yes |  |
| CMV-VSV-L | DPLA3191 | 132 | No, ECC fails | No VSV-L |
| CMV-VSV-L | DPLA3191 | 133 | No, ECC fails | No VSV-L |
| CMV-VSV-G | DPLA3192 | 134 | No, ECC fails | No VSV-G |
| CMV-VSV-G | DPLA3192 | 135 | Yes |  |

Being able to differentiate plasmids by their serial number makes it possible to associate different sets of documents with plasmids sent to different users. Just like the VIN allows multiple cars of the same make and model to be associated with different titles and insurance policies, it is desirable to be able to customize the sets of documents associated with a plasmid sent to a specific user. At the very least, each plasmid exchange or transaction is ruled by a unique agreement. In some cases, users may only need a user manual, while others may need access to the annotated plasmid sequence. By serializing self-documenting plasmids, it becomes possible to link each variant to a URL pointing to a folder, allowing the plasmid developer to share custom information with the individual recipient of a uniquely identifiable plasmid.

Even though the 24 original plasmids were not related to one another, we could find a conserved sequence between the origin of replication and the ampicillin resistance gene. This allowed us to insert the digital certificate in the same location for all of the plasmids. Inserting the certificate in the plasmid backbone minimized the risk that the certificate might interfere with the insert function. In addition, this greatly simplified the cloning of the certificate in the plasmid. We designed a pair of divergent primers positioned around the certificate insertion point and linearized all the plasmids by inverse PCR. The self-documenting plasmids were assembled by homology-based ligation of the linearized plasmids with synthetic DNA fragments encoding the digital certificate. For each ligation, we sequenced two colonies and assembled plasmid sequences using long and short read technologies as previously described [34,35].

#### Embedded documentation

We illustrate this self-documentation strategy using one of the 20 plasmids that we certified, a 4,966 bp mammalian expression vector expressing the GFP gene under the control of the CMV promoter, DPLA3122-CMV-mEGFP (Figure 3). We encode two types of text files as common examples of plasmid documentation. We designed two plasmids that encode GenBank files containing sequence annotations and a third that encodes a text file containing the Open MTA, a standardized document that can be used to quickly establish global sharing agreements (available at https://biobricks.org/open-material-transfer-agreement/) [36].

When possible, the detail in documentation should be maximized. However, certificate size scales with the size of the file and larger plasmids are generally harder to transform or transfect into cells [37-39]. We therefore created two versions of a GenBank file. DPLA3122-CMV-mEGFP_detailed.gb is 11 kilobytes (KB) and includes 18 features with detailed descriptions, including fields like notes and amino acid coding sequences, as well as some information about the source. The annotations in the GenBank file are encoded in a feature that is 6,804 bp long, resulting in a 12,027 bp self-documenting plasmid (DPLA3122-detailed-PID171.gb) (Figure 3). DPLA3122-CMV-mEGFP_summary.gb is a simplified version of the GenBank file that is only 8 KB. The sequence and features are the same as in the previous file, however, the file preamble has been removed and each feature only has a label and no other field. Having fewer annotations in the GenBank file reduces the size of the feature encoding the sequence annotations to only 2,064 bp. (Figure 3). The Open MTA text file was about 8 KB and was converted into an annotated ‘openmta.txt’ feature that was 11,218 bp, leading to the final 16,441 bp plasmid (DPLA3122-file-PID192). The certificates were ligated into the same place in the linearized plasmid backbone as above.

### Retrieving the documentation

#### Documentation reference

The documentation for certified plasmids can be accessed directly from *de novo* assembly FASTA files (Figure 2). The author ID found in the GSIN certificate is used as a public key to verify the data stored in the hashed signature. If needed, errors in the plasmid sequence are corrected using the error correction code (ECC). The plasmid ID is used to access the link to documentation stored in the database.

We obtained at least one clone matching the self-documenting plasmid sequence with complete sequence accuracy for 24 of the 48 plasmids (50% success rate) (Table 1). In two samples, 1 or 2 mismatch mutations were identified and fixed by the ECC (54% verification rate). However, the ECC fails to fix insertion or deletion mutations, and for 4 of the 24 plasmids, we could not produce a self-documenting variant. Reanalysis of several failed samples using an additional bacterial clone showed successful certification, indicating that failure resulted from factors in the ligation and assembly processes rather than an inability to certify certain plasmids. Indeed, 9 plasmids showed unexpected insertions seemingly related to primer dimer formation during the ligation process, accounting for about 40% of the failed certifications (Table 1).

For each sample that passes verification, with or without error correction, the plasmid’s identity, origin, and a URL to its documentation is revealed (Figure S2). Each plasmid variant with a different serial number can be linked to personalized documentation depending on the terms of use and sharing policies set by the developer. The URLs in our certified plasmids linked to fully annotated GenBank files, but one can imagine leaving certain sequences unannotated to protect proprietary sequence information. Thus, users of certified plasmids can get the information needed to act responsibly and in line with the terms set forth in the documentation without prior knowledge of the plasmid and without direct contact with the plasmid developer. Yet, the developer can be easily contacted with questions or concerns, especially when plasmids are error corrected. Furthermore, unauthorized or misuse of certified plasmids can be traced between developers and users.

#### Embedded documentation

We obtained at least one clone for each of the three self-documenting plasmids that verified successfully. The same information embedded in the standard GSIN data blocks is revealed as above, with the addition of the embedded files. All GenBank and text files were obtained with no errors. While the URL can still be used to share documents, all the pertinent information about a plasmid is directly embedded and accessible with no prior knowledge. This technology links the previously disparate molecule and documents into one system that can be easily verified throughout the plasmid’s lifecycle of synthesis, sharing, propagation, experimentation, and storage.

### Functional Characterization

Practical implementation of the GSIN technology requires the certificates to be biologically innocuous. We previously showed that the 512 bp certificate had no effect on bacterial colony growth or lacZ expression [31]. Here, we further characterize the effects of certification in both bacterial and mammalian cells. We first compared the yield of plasmid DNA extracted between the 20 plasmids successfully certified with the 164 bp signature and their uncertified counterparts and found no significant difference (Figure 4). Furthermore, we used spectral flow cytometry to compare EGFP expression in the five different versions of DPLA3122-CMV-mEGFP: uncertified, certified, both GenBank-encoding plasmids, and the file storage plasmid. We transfected 1 µg of each plasmid solution into HEK293-T17 cells. There was no GFP expression in our unlabeled negative control group. We performed Dunnett’s test to compare each of our sample groups to the unsigned reference group (Figure 5). In terms of the percentage of the cell population that expresses GFP, most certified samples were not significantly different than the unsigned group (∼46%). However, samples transfected with Plasmid #192 encoding the text file had significantly smaller average GFP positive populations, around 30% (Figure 5a). Replication of this experiment resulted in the same statistical trends (data not shown).

**Figure 4.**
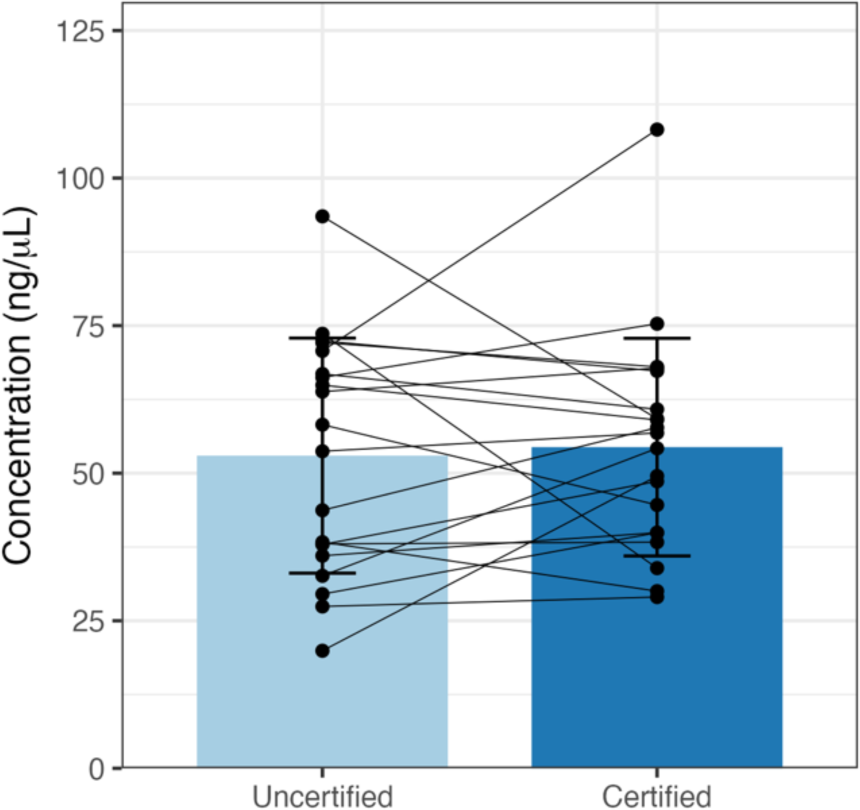
Insertion of certificate sequence does not affect plasmid yield. Yields from miniprep extractions of certi2ied plasmids and their uncerti2ied counterparts (connected by lines) were not signi2icantly different from one another (paired t-test, t(19) = −0.35, p=0.73). Data presented as mean ± SD.

**Figure 5.**
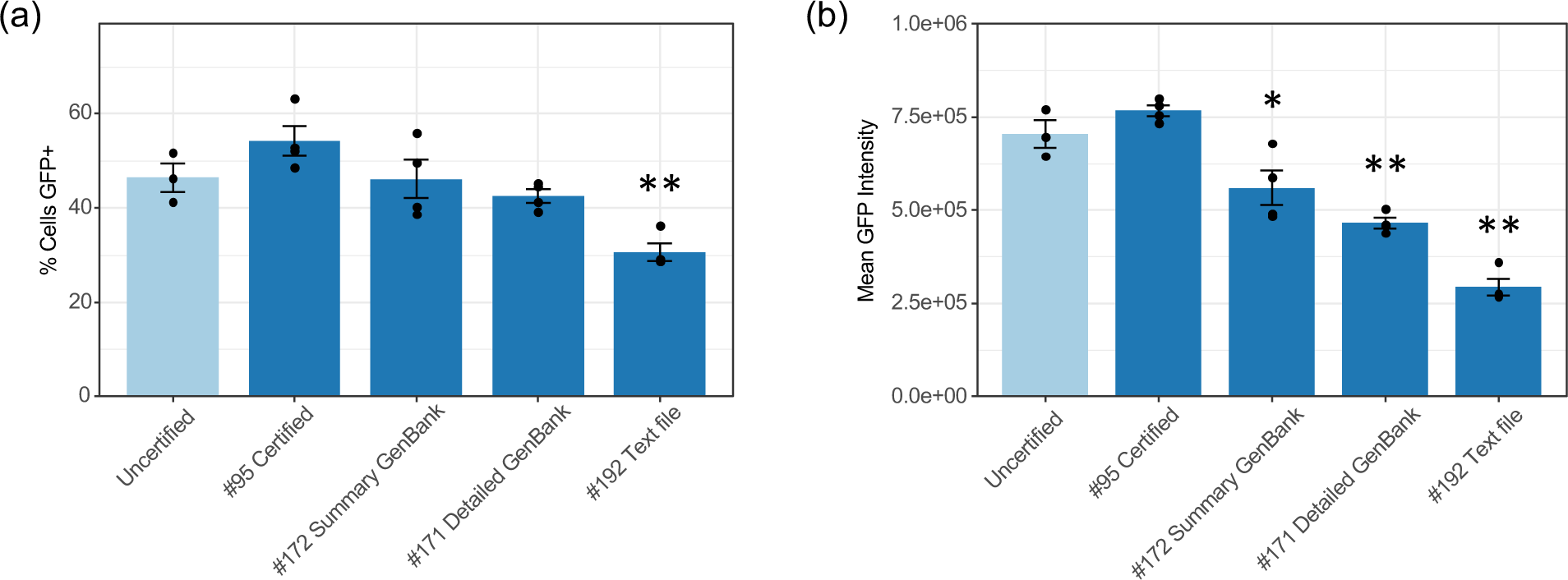
Functional protein expression is maintained after certificate insertion. Each certi2icate type was inserted into a CMV-mEGFP plasmid backbone. (a) The percentage of the cell population that expresses GFP decreases as plasmid length increases, with the 16 kb text 2ile-encoding plasmid being statistically different than the unsigned reference group. (b) The mean intensity of GFP also decreases as plasmid length increases, with statistically signi2icant differences in the GenBank- and text 2ile-encoding plasmids compared to the unsigned reference group. Data were analyzed using Dunnett’s test where * indicates p<0.05 and ** indicates p<0.01.

The mean GFP intensity in the certified CMV-mEGFP group was not significantly different from the unsigned reference group, whereas the intensity decreased as the size of the plasmid increased (Figure 5b). The sizes of the transfected plasmids ranged from 4,966 to 16,441 bp with decreasing GFP positive cell populations and GFP intensity in larger plasmids. Performing this experiment with equimolar concentrations of the plasmids provided the same results as the equimass experiments reported here (data not shown). These findings are consistent with studies that show that larger plasmids not only have lower transfection efficiencies, but that fluorescent protein expression (positive population size and intensity) decreases with larger plasmids, even in equimolar experiments [40,41]. It is thought that despite equivalent entry into the cell across plasmid lengths, differences in nuclear translocation and transcription underlie the relationship between plasmid length and gene expression [42]. Nevertheless, functional protein expression is maintained even when an 11 kb certificate is inserted in the plasmid.

## Discussion

The ability to track and verify plasmids throughout their lifecycle is critical to the security of the bioeconomy, yet the documentation of DNA sequences is often insufficient to verify their accuracy. GSIN certificates transform the plasmid sharing process by simplifying sequence verification performed by plasmid users and allowing developers to share customized documentation that is unique to each serialized plasmid. This simultaneously enables increased transparency in plasmid sharing as well as a novel approach to protecting proprietary sequence information. A major advantage of GSIN certificates over previous iterations of DNA watermarks is the implementation of a public-key cryptographic approach [19]. It assumes that the verifier is following a *de novo* plasmid assembly protocol and requires no knowledge of the reference sequence nor any specific primers to retrieve the embedded documentation. The GSIN workflow eliminates the need to evaluate each plasmid assembly against a reference sequence and manually catalogue errors, which accelerates screening experiments to produce accurate samples faster. Certified plasmids can be distributed, verified, and used for experimentation without relying on assumptions about the plasmid identity and the separate exchange of email attachments documenting the plasmid.

We demonstrate that certificates can be embedded into diverse plasmids and used to share documentation that is either hosted on a server or embedded directly into the plasmid. We show that insertion of the 164 bp signature has no effect on DNA yield and that insertion of the larger embedded file certificates does not inhibit functional protein expression in mammalian cells, which is consistent with findings from the first iteration of 512 bp certificates that had no effect on bacterial growth or lacZ expression [31].

A few factors contributed to our verification rate of only 54%. We encountered issues with the formation of primer dimers during the linearization of the plasmid backbone that, in some cases, led to large insertions of repeated primer sequence before or after the certificate sequence. Since we set out to linearize all plasmids in a conserved location in the backbone, this limited our options for primer design, leading to higher complementarity between the two primers than would be ideal. Choosing multiple colonies per sample, which is recommended to detect errors appearing only in some bacterial colonies [43], would have further improved our assembly rate. Despite errors during the plasmid assembly process, most plasmids were successfully verified and the software correctly differentiated between accurate and inaccurate plasmids.

The choice between the two strategies for developing self-documenting plasmids depends on the documentation that needs to be shared. The plasmids with embedded files have a secure, direct link between the molecule and the documentation. Practical use of these file storage certificates is currently limited to relatively small text files, as plasmids with larger files (images, pdfs, etc.) become difficult to synthesize and work with in the lab. Yet the 164 bp signature is sufficient for making a self-documenting plasmid. If needed, other file types, large files, and documentation datasets containing multiple files can be reliably shared using plasmids with the 164 bp signature scheme via a link to the documentation hosted online. Sharing through a URL can also allow the use of additional security features like user permissions and password protection. In future iterations of the GSIN technology, improved data compression techniques may be applied to increase the data payload to sequence length ratio [44]. Alternatively, we foresee this technology being expanded to allow for certifying smaller fragments within a plasmid, or for certifying gene networks composed of multiple plasmids, which could disperse large data files into multiple data blocks. Regardless, the examples presented here confirm that some of the most common types of documentation can be easily embedded in and retrieved from plasmids.

The GSIN workflow is available for academic use via a simple web interface (https://sign.genofab.com/). Here, we validate the GSIN technology and show that it is ready for implementation outside our laboratory. While there are no barriers to using the application in its current form, we acknowledge there are several avenues to extend the capabilities of the technology. The primary limitation of the technology in its current form is the inability to correct insertion or deletion (indel) mutations. This is especially important to individuals relying on nanopore-based DNA sequencing techniques, which are more prone to having indels in their assemblies [6]. Indels are more complicated than substitutions to correct, although some methods can do so [45,46]. However, the choice of an ECC involves a tradeoff between the scale of the errors being corrected and the length of the code sequence. In some cases, mutations render a plasmid unacceptable and so the ECC can be minimized; in others, it may be worthwhile to have a large ECC that allows a greater number of mutations to be corrected. Future updates could offer multiple error correction schemes after further experimental validation [47].

Screening procedures for certificate sequences and embedded files could also be implemented. Here, we used the Basic Local Alignment Search Tool (BLAST) by NCBI to confirm that none of the GSIN sequences had significant similarity to any sequence in the nucleotide collection [48,49]. In future iterations, the certification software could implement a screening algorithm to detect and correct problematic sequences within the certificate sequence. This could include sequences with similarity to existing genes, sequences of concern, as well as novel genetic parts (e.g., new start/stop codons, cryptic promoters, homopolymeric sequences, etc.). This is especially important considering it has already been shown that malicious software (“malware”) can be encoded in DNA and unleashed onto the computer that is sequencing the strand of DNA [50]. Files encoding malware or computer viruses could theoretically be embedded in a similar scheme as we used for embedding GenBank and flat text files. As is, the onus is on the plasmid designer to screen their sequences and adhere to not only biological constraints but federal regulations surrounding DNA sequences before synthesizing the novel construct [51]. This points to an unmet need for approaches for determining whether new synthetic sequences are innocuous or potentially harmful on their own or in tandem with other constructs.

## Concluding Remarks

GSIN certificates transform the plasmid documentation process by directly associating a plasmid molecule with its digital documentation, either as a link to data stored on a server or a data file embedded into the sequence itself. This system simplifies sequence verification and allows customized documentation sharing by developers, providing an avenue for the protection of IP throughout the plasmid distribution process. Plasmid users can easily sequence verify certified plasmids, determine their origins, and obtain the documentation needed to use the plasmid according to the terms of use set by the developer, all without any prior knowledge of the plasmid sequence. We show that certificates do not adversely affect DNA yield or mammalian gene expression, although further testing is needed to evaluate the practicality of inserting certificates into higher organisms (see Outstanding Questions). Widescale adoption of the GSIN will accelerate plasmid production pipelines, allow the validity of biological products to be frequently confirmed and improve biosurveillance, for example, by simplifying the identification of GMOs.

## Acknowledgements

This work was supported by the National Science Foundation #MCB-2123367 to J.P.; the National Institutes of Health R01GM147816 and R21AI168482 to J.P.; and a generous gift from the Suzanne and Walter Scott Foundation. Figure 2 was created in BioRender (https://BioRender.com/w02n093).

## Author Contributions

Conceptualization, J.P., S.P., C.-T.B.; Software, S.P.; Investigation, S.H.; Formal Analysis, S.H., C.- T.B.; Validation, S.H., C.-T.B., J.P.; Writing – Original Draft, C.-T.B.; Writing – Review & Editing, C.-T.B., S.H., S.P., J.P.; Supervision, J.P., C.-T.B.

## Outstanding Questions

▪ What is the ideal tradeoff between error correcting capability and error correction code length?
▪ What is the limit on file size that can be practically embedded into a plasmid?
▪ What challenges will arise in inserting and extracting certificates from higher organisms?
▪ Can certificates be expanded to verify sequences across multiple DNA molecules such as gene networks and genomes?
▪ Can improved binary to DNA encoding schemes, developed for DNA data storage, be used to streamline certificate size?
▪ What new cybersecurity risks are users of this technology exposed to? Can malware be inserted and shared with plasmids?

### Technology Readiness Box

We present a practical solution to link DNA molecules and their digital documentation. We experimentally validate certificates that make plasmids self-documenting by providing a link to their documentation or embedding the documentation in the DNA sequence. Ongoing efforts aim to evaluate commercial applications, and academic users can test this technology using a publicly available website. This technology has moved into the deployment phase corresponding to Technology Readiness Level (TRL) 7. Further experimental testing is needed to validate the technology beyond cell culture, but based on our results, we anticipate little to no adverse effect of certification on higher organisms. Improved data encoding techniques are still needed to increase the practicality of embedding large data files into certificates of reasonable length. Areas for immediate future development include extended error correction capability, enhanced user permission settings, and the integration of screening algorithms for biocompatibility and sequences of concern. Certification improves the documentation process for plasmid sharing with implications for basic research, bioproduction, biosurveillance, clinical and commercial development.

## Methods

### Certification Software Development

The GSIN web application is publicly available at https://sign.genofab.com/. It was developed with Java using Apache Tomcat on the server-side and React.js on the client-side. Firebase Firestore, a non-relational database service from Google Cloud, is used for backend storage. Firebase is also used for user authentication using email and password credentials. The app is deployed on an Amazon Web Services Elastic Beanstalk instance. Whenever a new plasmid is generated in the app, an entry in the database is created. This entry contains the name of the new plasmid, a backup of the GenBank File as text, the ID of the user who generated the file, and the URL.

The algorithm used for certificate generation and verification is as previously published [31], with modifications to how user IDs are generated and used as private keys. The previously published method signed each plasmid using a private key derived from the signer’s unique ORCID, necessitating the user to have an ORCID account. Instead, this new software assigns a unique user identifier (uuid) to each account upon signing up. The uuid is auto generated by the Firebase authentication service, but it does not conform to same 16-digit number format as the ORCID. The Firebase uuid is converted to the ORCID format using a truncated SHA-256 hash. Each users’ private key is now derived from the truncated hash instead of the ORCID.

Files embedded within the signature are compressed using the zlib library implementation of the DEFLATE algorithm, a lossless data compression algorithm. The compression ratio is variable, depending on the data type and repetition of the structure.

### Experimental Validation of Certified Plasmids

#### Reagents

The Zyppy-96 Plasmid MagBead Miniprep Kit was purchased from Zymo Research (Irving, CA, USA, #D4102). The Oxford Nanopore Rapid Barcoding Kit and R10.4 flow cell were purchased from VWR (USA, #SQK-RBK114.96 and #FLO-MIN114). For the short-reads, the MiSeq Nano V2 cartridges were purchased from Illumina (San Diego, CA, USA. #20060059 and #MS-103-1003). Library preparation kits (ExpressPlex™ Formula M) for the short-reads were sourced from SeqWell (Beverly, MA, USA).

#### Biological Resources

The plasmids used for library preparation, sequencing, and analysis were obtained from two vendors or were designed and manufactured in house. Eleven plasmids were synthesized and sequence-verified by Twist Biosciences (San Francisco, CA), and five were procured from Addgene (Watertown, MA, USA). The remaining three plasmids were made using fragments and vectors sourced from Twist Biosciences (San Francisco, CA). The array of plasmids was given unique identifiers (e.g., DPLA1234). Primers for plasmid linearization were sourced through Integrated DNA Technologies (Coralville, IA, USA).

#### Plasmid Construction

To certify the plasmids used in this paper, two approaches were used in plasmid construction. For the smaller signatures, uncertified plasmids were first linearized via PCR amplification using the NEB Q5 Hot Start High Fidelity 2X Master Mix (Ipswich, MA, USA. #M0494S) according to the manufacturer’s instructions with 25 cycles and annealing temperatures determined using the NEB Tm calculator tool (tmcalculator.neb.com/). Post linearization, plasmids were quantified using both a Synergy HTX plate reader for A260/280 measurements and a TapeStation 4200 using the genomic screen tapes and reagents to verify fragment size, purity, and quantity (Santa Clara, CA, #5067-5365 and #5067-5366). All certificate sequences were ordered as synthesized gene fragments, each with 25-30 bp of overlap to facilitate ligation into the plasmid backbone. Plasmid assembly was then performed from fragments and the linear plasmids, using the NEBuilder HiFi DNA Assembly Master Mix (Ipswich, MA, USA., #E2621L) according to the NEBuilder design tool available on the manufacturer’s website (with the exception of an increased incubation time to one hour). In brief, equal molar equivalents of fragments were added to a tube with the master mix. For the larger signatures, (e.g., the embedded file and documentation), plasmid construction consisted entirely of fragments sourced from Twist Biosciences (San Francisco, CA), with the same ligation process described above. The ligation solution for each plasmid was transformed into chemically competent *E. coli* cells and grown overnight.

#### Plasmid Minipreparation

The plasmid DNA pre- and post-assembly was extracted from each of the 21 isolates on an epMotion 5075 TC liquid handler (Eppendorf, Hamburg, DE) using the Zyppy-96 Plasmid MagBead Miniprep Kit (Zymo Research, Irving CA, USA), according to manufacturer’s instructions with the following modifications: pipet mixing during the lysis and neutralization steps, increased Zyppy wash buffer to 800 uL, and an extended elution time of 10 minutes. Samples were all quantified on a Synergy HTX plate reader to determine the quality and quantity of the solutions after purification. Statistical analysis of the DNA yield was performed using R.

#### Plasmid Isolation and Sequencing

The plasmids were sequenced with the Oxford Nanopore Rapid Barcoding Kit V14.96 according to the manufacturer’s specifications using 50 ng per plasmid. The samples were set to run for 24 hours on the MinION with the following parameters: super accurate basecalling, barcode trimming, and basecalling every hour. The Express Plex Formula M kit was used to prepare the plasmids for short read sequencing according to the manufacturer specifications using 2.5 ng per plasmid. The samples were loaded on to the Illumina MiSeq and demultiplexed to reflect the indexes supplied within the kit. All data was processed into final assemblies using PlasCAT, a hybrid assembly pipeline that takes both Illumina short-reads and Nanopore long-reads as input to generate assembly sequences [52].

#### Transfections and Flow Cytometry

Plasmids were transfected into HEK 293T/17 cells using Lipofectamine 2000 (Invitrogen, ThermoFisher, MA, USA) according to the manufacturer’s instructions in a 24-well plate. Cells were transfected at around 70% confluency and the growth media (DMEM supplemented with 10% v/v fetal bovine serum) was changed 4 hours post-transfection. After 24 hours of growth, cells were harvested using TrypLE Express and phosphate buffered saline according to the manufacturer’s instructions adjusting the volume for the 24-well plate (ThermoFisher, MA, USA). EGFP expression was quantified using the Northern Lights spectral flow cytometer (Cytek, Fremont, CA) to measure 10,000 events per sample. Unmixing was performed with autofluorescence extraction, using unstained cells as a control. Statistical analysis of the spectral data collected was performed using R.

#### Data Availability

All data files generated for this study are available at: https://figshare.com/s/fbc52c8931ad07a3c83a. This includes the embedded files (DPLA3122-CMV-mEGFP_detailed.gb, DPLA3122-CMV-mEGFP_summary.gb, and openmta.txt), GenBank files of certified plasmid designs, FASTA sequencing assembly files, miniprep and flow cytometry data, and R markdown files.

