## Supplemental Figures for "Self-Documenting Plasmids"

### Self-Documenting Plasmids: Supplementary Figures

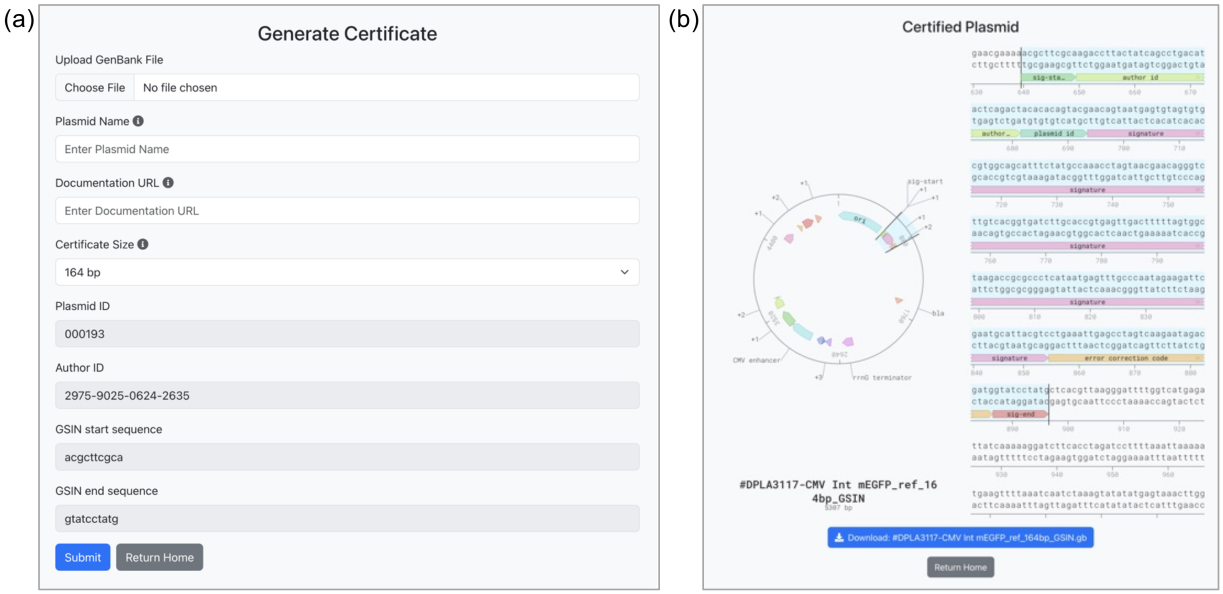


**Figure S1. Plasmids are certified through a simple web application. (a)** To generate a certificate, users upload a GenBank file to the form, then designate a plasmid name, URL, and certificate size (164 bp signature, 512 bp signature, annotations, file storage). Plasmid and author IDs are automatically generated by the application, and delimiter (start/end) sequences are standardized. Identical plasmids can be certified with unique plasmid IDs and URLs. A plasmid map viewer then appears for the certifier to select a location to insert the certificate. URLs are not embedded in the sequence, so can be changed by the certifier later using the 'Manage Plasmids' feature of the application. For file storage plasmids, an entry field appears for the user to upload a file. **(b)** A GenBank file with the certificate features annotated is available to download. Any part of the plasmid sequence can be easily selected (blue highlight) and copied.

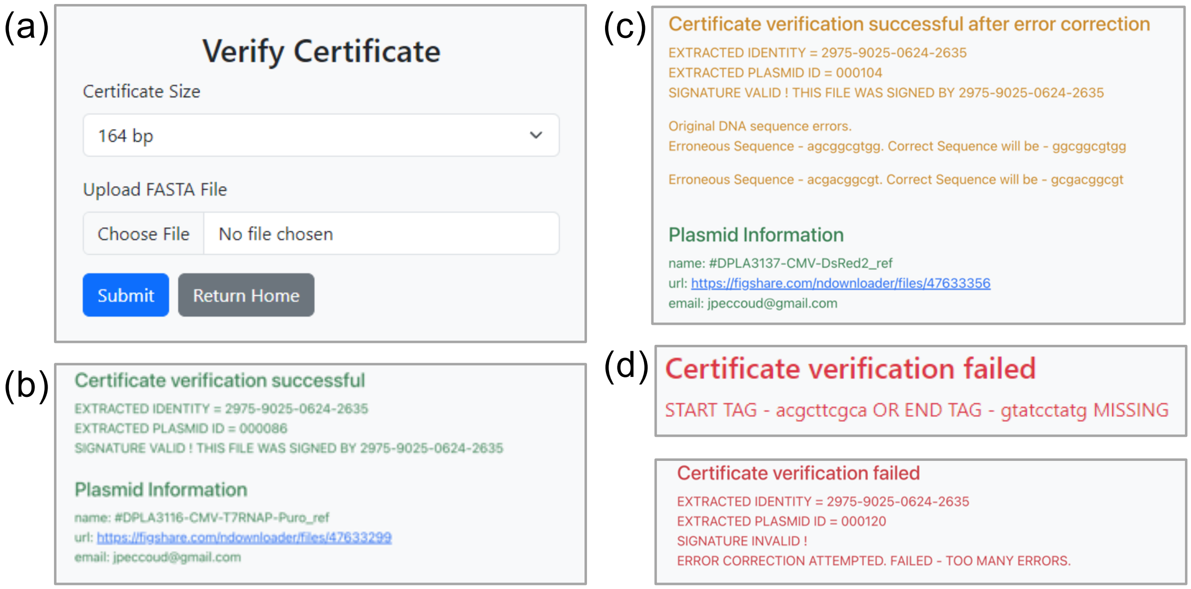


**Figure S2. Certificate verification is accurate and secure. (a)** Verification of a certified plasmid requires only the type of certificate expected and a FASTA file of the plasmid assembly. **(b)** If the assembly matches the expected sequence, plasmid and author information is revealed, including a URL to documentation about the plasmid, in this case, a GenBank file hosted on figshare. Files from plasmids with embedded documentation will be automatically downloaded. **(c)** The error correction code can identify and accept up to two mismatches in the certified plasmid sequence to successfully verify a plasmid. **(d)** Plasmids without signature delimiters (top), more than two mismatches or any insertions or deletions (bottom) will fail verification.
